# *In vivo* selection reveals long non-coding RNAs implicated in colon to liver metastasis

**DOI:** 10.1101/2022.05.23.493110

**Authors:** Artin Soroosh, David M. Padua, Elizabeth Videlock, Diane Bui, Ami Patel, Charalabos Pothoulakis, Carl Robert Rankin

## Abstract

Colorectal cancer (CRC) is the third most common malignancy in both American men and women. Most of the deaths attributed to CRC are a result of metastatic spread to the liver. In this study, colon cancer cells that highly metastasized to liver *in vivo* were compared to less metastatic parental cells to investigate the role for long non-coding RNAs (lncRNAs) in CRC metastasis. The highly metastatic daughter cells (LS-3B) were found to be 63-fold more metastatic than the parental cell line (LS-PAR) *in vivo*. A lncRNA microarray comparing LS-PAR and LS-3B cells revealed that 104 lncRNAs had fold changes > 2.0 and an FDR < 0.05. Real time PCR mediated validation revealed many lncRNAs exhibited high fold changes such as a 60-fold increase in LOC101448202, a 20-fold increase in MRPL23-AS1 and 50-fold decreases in GNAS-AS1 and LOC101928131. *In vivo* metastasis differences could be recapitulated *in vitro* as LS-3B cells closed wounds faster than their parental LS-PAR cells. However, intestinal epithelial cancer cells with robust downregulation of MRPL23-AS1, C1QTNF1-AS1, GNAS-AS1, LINCR-0002 and LOC101448202 failed to display differences in comparison to controls in *in vitro* migration assays. Three of the five lncRNAs with microarray probes for currently available GEO-datasets were significantly altered in liver CRC-associated tumor biopsies as compared to the primary tumor of non-metastatic CRC. Further studies on the lncRNAs identified will better define their roles in metastasis and how they might be useful if targeted therapeutically.

## Introduction

While colorectal cancer (CRC) rates in the US have declined over the past twenty years (3), in 2019 over 100,000 people were diagnosed with CRC even with saturated surveillance and pre-cancerous polyp removal. CRC is still the third leading cause of death in the US, causing an estimated 50,000 deaths in 2019 (42). Primary colonic tumor growth is rarely the cause of death in most patients -- instead, mortality is dependent on the mobilization of the cancer to distant tissues and subsequent failure of the metastasized organ. Colon cancer cells target the liver in over 70% of metastases by utilizing complex mechanisms to travel from the primary tumor site to distant targets (2). Primary tumor resection followed by chemotherapy is the first line of treatment to local CRC, but metastatic CRC requires prolonged chemotherapy. Unfortunately, median survival rates remain at less than 2 years for patients with metastatic CRC (47). Given the high morbidity and mortality of metastatic CRC, new therapeutic targets are paramount in improving outcomes for this disease.

A new class of potential therapeutic targets for CRC metastasis are long non-coding RNAs (lncRNAs), as over the past 10 years the number of well annotated lncRNAs has doubled. These genes produce non-protein coding transcripts greater than 200 base-pairs and have been shown to participate in most cellular functions (20). As of 2019, over of 16,000 long non-coding RNAs have been annotated in humans (49). While a few lncRNAs have been identified to be metastatic drivers, namely H19 (31), MALAT1 (17), and HOTAIR (23), most alterations found in lncRNAs of *in vivo*-selected metastatic cells have not been profiled. *In vivo* selection relies on the fact that in a pool of mildly metastatic cancer cells, some will mutate over time to aggressively metastasize to the liver (32). These mutated cells have been shown to have major alterations in miRNAs and mRNAs, enhancing pro-metastasis genes and reducing anti-metastasis genes (6, 22, 44).

Here, *in vivo* selection and *in vitro* migration studies were used to identify and characterize lncRNAs important for CRC metastasis. After validating that the *in vivo* selected daughter cell line (LS-3B) was greater than 60-fold more metastatic than the parental cell line (LS-PAR) a microarray was performed (30). Many lncRNAs displayed high fold changes between the highly metastatic cells and mildly metastatic cells. Many of the lncRNAs identified were in the genomic vicinity of pro-metastatic protein-coding genes. Studies using RNA interference followed by *in vitro* metastasis assays failed to identify functional differences in cells with down-regulated lncRNAs. However, lncRNA analysis comparing CRC patients that never developed liver metastasis to CRC patients with liver metastasis mirrored what was observed in the *in vivo* selected cells.

## Materials and Methods

### Cell culture

The LS-147T derivative cell lines, LS-Par and LS-3B were a gift from the Tavazoie Lab. The SW480 cells used were purchased from American Type Culture Collection (Manassas, VA). All three cell lines were cultured in DMEM (Corning, Corning, NY) supplemented with 10% fetal bovine serum (Sigma-Aldrich, St. Louis, MO).

### *In vivo* metastasis assay

The *in vivo* metastasis assay was performed at the West Los Angeles Veterans Administration under the IACUC #01003-17. Seven-week old *Prkdc*^*scid*^ mice were used (Jackson Laboratory, Bar Harbor, ME). Mice were anesthetized with isoflurane and 20 µg of buprenorphine-SR was administered per mouse. After shaving and betadine swabbing, a small incision was made in the left subcostal region to access the spleen. 0.5 million cells suspended in 50 µl of saline was injected into the spleen through a 26 gauge needle while the spleen was held with forceps. Almost immediately after injection the spleen was then removed by thermal cautery. To analyze the luciferase signals, 100 µl of 15 mg/mL Luciferin (Gold Biotechnology, St. Louis, MO) was injected retro-orbitally. An IVIS imager was used to measure the luciferase signal (Perkin Elmer, Waltham, MA).

### Microarray

An Arraystar human lncRNA microarray version 4.0 was used for the global expression profiling of lncRNA and mRNA transcripts (Rockville, MD). The sample preparation and microarray hybridization were performed based on the manufacturer’s standard protocols with minor modifications. Each sample was amplified and transcribed into fluorescent cRNA along the entire length of the transcripts without 3’ bias utilizing a random priming method (Arraystar Flash RNA Labeling Kit, Arraystar). The labeled cRNAs were hybridized onto the Human LncRNA Array v4.0 (8 × 60K, Arraystar). After washing the slides, the arrays were scanned by the Agilent Scanner G2505C. Agilent Feature Extraction software (version 11.0.1.1) was used to analyze the acquired array images. The microarray data is deposited in NCBI under GEO# GSE145749.

### RNA isolation and qRT-PCR

An Aurum RNA mini kit was used to purify total RNA from the cells (Bio-Rad, Hercules, CA). For real time PCR experiments, an iScript reverse transcription kit was used to convert one microgram of RNA to cDNA (Bio-Rad, Hercules, CA). No reverse transcriptase controls were included. PCR was performed using iTaq universal SYBR green and a CFX connect 384 real time PCR machine (Bio-Rad, Hercules, CA). Primer-BLAST was used to design the primers (50). Primers were from Integrated DNA Technologies (Coralville, IA). Primer sequences are listed in Supplemental Table 1a. Supplemental material is available at https://doi.org/10.6084/m9.figshare.11879472.v1.

### Transfections *and in vitro* migration assays

To transfect SW480 cells, 50 nM LinCode LincRNA siRNA or negative control siRNA (Dharmacon, Lafayette, CO) was diluted in warm Opti-MEM (Gibco, Waltham, MA) and 11 µL/mL of Lipofectamine RNAiMax (Invitrogen, Carlsbad, CA). The siRNA sequences are provided in supplemental table 1b. Supplemental material is available at https://doi.org/10.6084/m9.figshare.11879472.v1. SW480 cells resuspended in transfection mix were seeded on 12-well plates and after 3-4 days were 80-90% confluent. Then, a sterile 10-uL pipette tip was used to gently scratch a thin wound gap down the middle of each well in one direction with the tip remaining perpendicular to the bottom surface of the well. After scratching, the cells were washed once with phosphate-buffered saline containing calcium and magnesium to remove any detached cells, and fresh media was added to each well. A section along each scratch wound was then marked. An EVOS XL Core Configured microscope (Advanced Microscopy Group, Bothell, WA) at 10x magnification was used to image the wound immediately after the scratch was made and 24 hours later. ImageJ software was used to quantify the area of the gap (41).

### Bioinformatics

GEO2R was used to analyze publicly available clinical microarray data comparing patients with primary CRC tumors from patients without metastatic tumors to CRC derived liver metastasis (4). The GEO accession number for PRKCQ-AS1 was GSE75050. The GEO accession number for LINC00635 LINC00842 and DLX6-AS1 was GSE95423.

### Statistics

Microarray data was quantile normalized and the GeneSpring GX v12.1 software package was used to determine statistical significance (Agilent Technologies, Santa Clara, CA). Quantitative real time PCR was used to determine relative transcript levels from the cycle threshold values and are expressed as fold change to the average control (40). HPRT1 was used as a housekeeping gene. For analysis between two groups, a Student’s t-test was used. Data was considered significant when p < 0.05. All group comparisons used Student’s t-tests and two-tailed p-values are reported. For microarray data, false discovery rate (FDR) was determined with the Benjamini-Hochberg method.

## Results

### Over 100 long non-coding RNAs are altered in highly metastatic cells compared to mildly metastatic cells

As a method to identify long non-coding RNAs (lncRNAs) involved in colon to liver metastasis, a set of previously developed and characterized mildly metastatic parental KRAS mutant colon adenocarcinoma cells, LS147T, and *in vivo* selected highly metastatic cells were used (30). To create highly metastatic daughter cells, the parental cells were first injected into the spleen (Figure 1a). The cells then go into the portal vein circulatory system and the spleen is then removed (Figure 1b). After 21 days, the livers are removed and cells which have mutated to preferentially metastasize to the liver can be analyzed or further purified to create more aggressive metastatic cells. As a proof of concept, the mild metastatic parental cells, LS-PAR, were compared to the highly metastatic *in vivo* selected LS-3B cells. It was observed that after 21 days only metastatic tumors developed in the liver and this only occurred in mice injected with LS-3B cells (Figure 1b).

**Figure 1:**
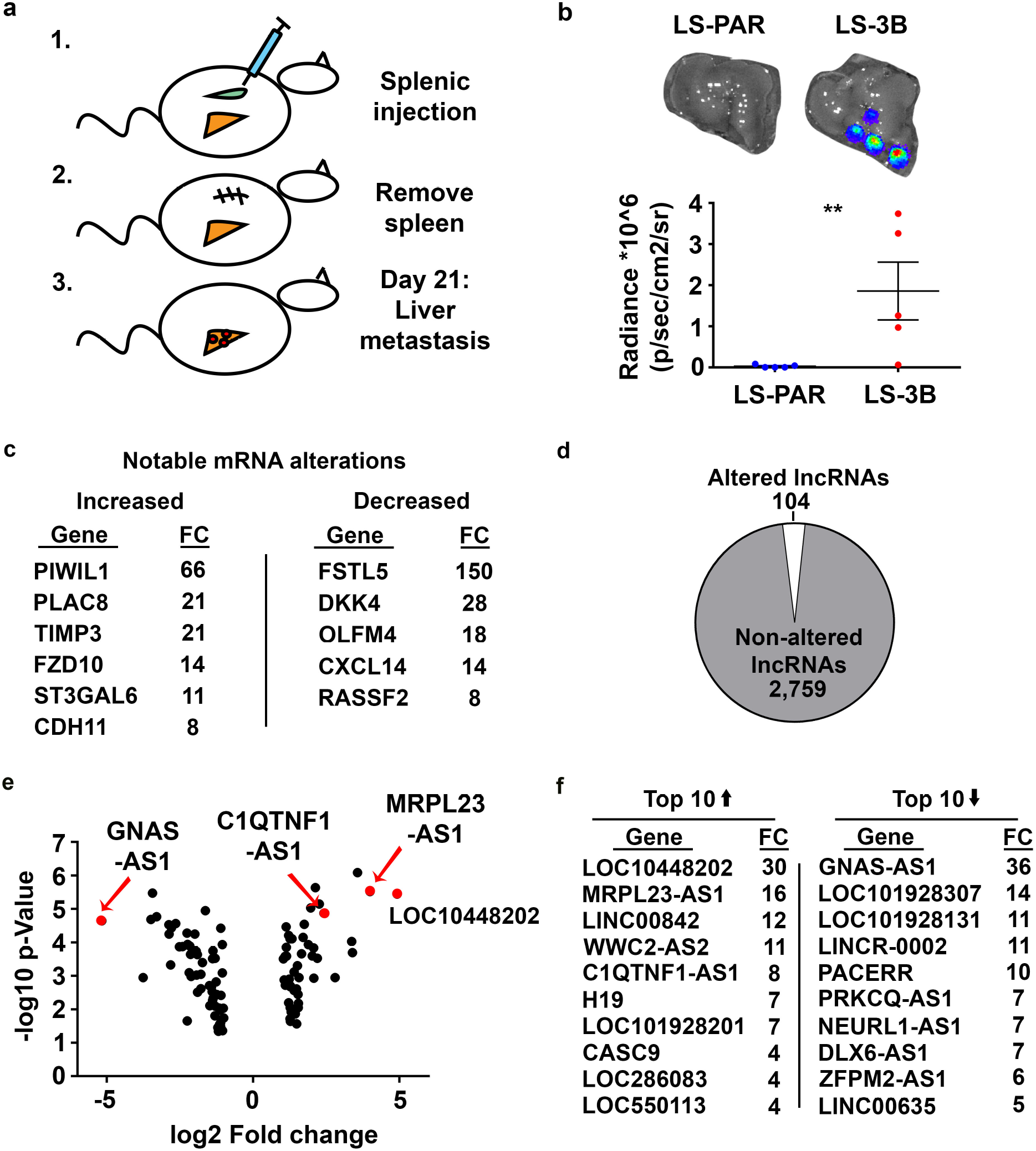
Over 100 long non-coding RNAs are altered in highly metastatic cells compared to mildly metastatic cells. (a) A cartoon for a model of CRC liver metastasis. (b) After 21 days of metastasis, mice were injected with luciferin, livers were removed and an IVIS imager was used to analyze tumor Radiance. n=5 per group, ** p <0.01. (c) A table listing well-known protein coding genes involved in metastasis that displayed high fold changes and p-values less than 0.001 in LS-3B cells compared to LS-PAR cells after microarray analysis. There were 3 independent samples per group in the microarray. (d) A pie chart comparing Ref-Seq lncRNAs that were observed to display a fold change > 2.0 and an FDR <0.05 to total Ref-Seq lncRNAs detected in the microarray. (e) A volcano plot comparing statistical significance (y-axis) to fold change (x-axis) for the 104 lncRNAs altered in LS-3B cells compared to LS-PAR cells. (f) A table listing the top 10 up and down-regulated lncRNAs in LS-3B cells compared to LS-PAR cells.

To identify mRNAs and lncRNAs that were altered in the highly metastatic LS-3B cells compared to mildly metastatic LS-PAR cells, a microarray was performed. While the purpose of the experiment was to analyze lncRNAs differences between these cells, alterations in mRNA were also assessed. The power of analyzing the transcriptome between parental and aggressive cancer cells is evident, as many previously identified clinically relevant metastatic protein coding genes were highly altered in LS-3B cells compared to LS-PAR cells (Figure 1c). *PIWIL1* and *FSTL5* were the most altered mRNAs between LS-3B and LS-PAR cells, with a 60-fold increase and a 150 decrease, respectively (9, 51). Accordingly, genes encoding the secreted anti-metastasis proteins, DKK4 and CXCL14, were 28 and 14 fold decreased in LS-3B cells compared to LS-PAR cells (13, 16). The pro-metastasis mRNAs *PLAC8, TIMP3, FZD10, ST3GAL6* and *CDH11* were 8 to 21 fold increased in the highly metastatic cells compared to their parental counterparts (14, 15, 19, 26, 28).

When selecting for lncRNAs in the RefSeq database (33) that had a FDR less than 0.05 and a fold change greater than 2, 104 lncRNAs were identified to be altered out of 2,863 total lncRNAs expressed in LS-3B compared to LS-PAR cells (Figure 1d). There were 47 elevated and 57 decreased lncRNAs in LS-3B cells compared to LS-PAR cells. A volcano plot of this data, displaying statistical significance versus fold change for lncRNAs in LS-3B cells versus LS-PAR cells, highlights the major alterations to lncRNAs in highly metastatic cells (Figure 1e). As differential expression of most lncRNAs was highly statistically significant, these genes were categorized by fold change (Figure 1f). While most of the mRNAs seen to be altered in metastatic cells have been mechanistically analyzed, of the top 20 lncRNAs up and down-regulated in metastatic cells only *H19, CASC9, GNAS-AS1, LINC00842* and *DLX6-AS1* have been studied in the context of cancer as observed by PubMed searching.

### An independent method verified the alterations to lncRNAs in highly metastatic cells

As the quality of high-throughput assays can vary between experiments, real time PCR was used for validation. Based on literature searches and methodological (primer design) considerations, 13 genes (6 of the top 10 elevated and 7 of the top 10 decreased lncRNAs in LS-3B cells compared to LS-PAR cells) were selected for validation. All of the genes measured by real time PCR were significantly altered (6 with p < 0.001, Figure 2). Notably, *LOC101448202* was 60-fold elevated and *GNAS-AS1* was 50-fold decreased in LS-3B compared to LS-PAR cells. Now validated, these results represent new therapeutic opportunities for the treatment of metastatic cancer.

**Figure 2:**
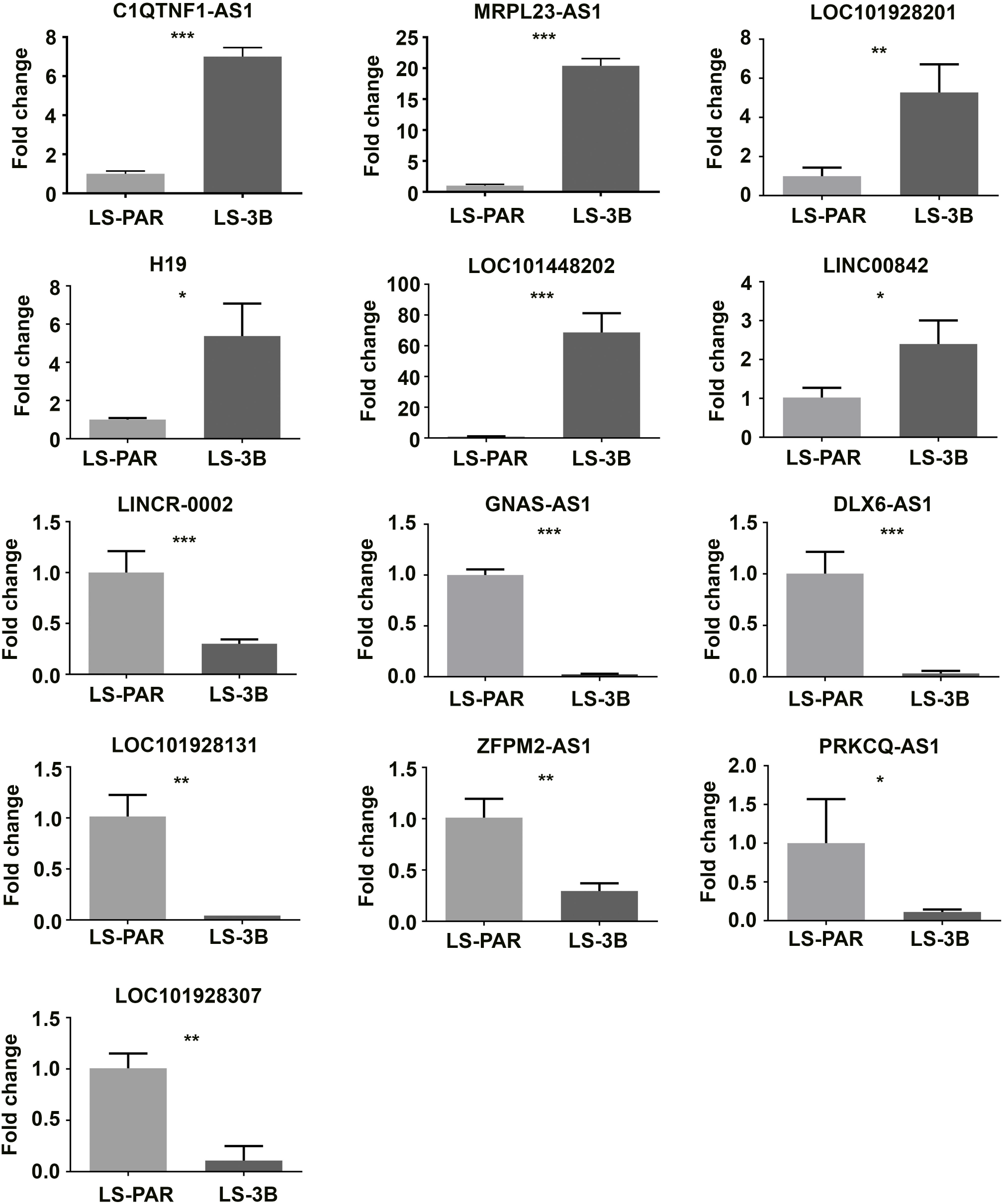
An independent method verified the alterations to lncRNAs in highly metastatic cells. Real-time PCR was used to measure the levels of lncRNAs in LS-PAR and LS-3B cells. There were 3 independent samples per group. * p <0.05, ** p <0.01, *** p <0.001.

### lncRNAs identified by *in vivo* selection are also altered in clinical samples and reside near metastasis associated protein coding genes

Most of the genes identified in the microarray had never been studied nor had been assigned names. As many lncRNAs have been shown to function by regulating gene expression in *cis* (45), to better understand their possible functions in metastasis an analysis of their genomic surroundings was performed next. *LOC101448202* overlaps with Collagen 5A1 (*COL5A1*) gene and is near Ficolin-2 (*FCN2*) (Figure 3a), and *COL5A1* and *FCN2* have both been associated with metastasis (10, 48). However, cells treated with siRNAs against *LOC101448202* failed to display differences in *COL5A1* expression (data not shown). When comparing the nuclear a cytoplasmic localization of LOC202, it is entirely in the nucleus. *LOC101928307* is upstream of the p53 and angiogenesis associated gene *ADGRB3* (Figure 3b) (11). *LINCR-0002* overlaps with and is adjacent to two mRNAs that both regulate NFkB signaling (5, 21), which is important for metastasis (Figure 3c) (39). While *LINC00842* is flanked by 3 protein coding genes: *FAM25BP, ANXA8L1* and *NPY4R*, none of the nearby genes have been identified in cancer (Figure 3d). *GNAS-AS1* is upstream from the metastasis associated microRNA MIR296 (Figure 3e) (25). It is likely that for some of the lncRNAs identified to be altered in metastatic cells they function through regulating transcription of nearby protein coding genes.

**Figure 3:**
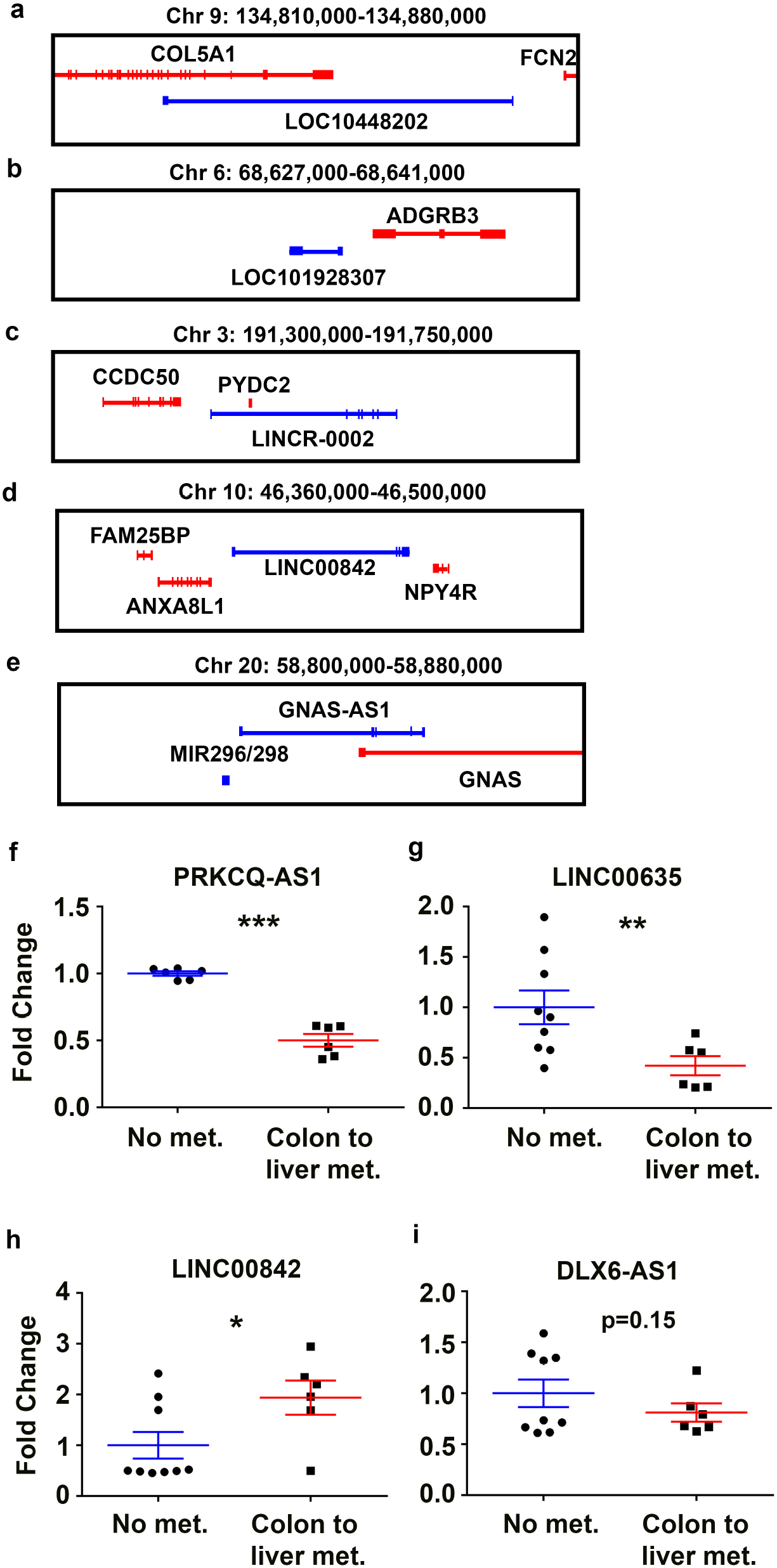
lncRNAs identified by *in vivo* selection are also altered in clinical samples and reside near metastasis associated protein coding genes. Screenshots of the UCSC genome browser (18) for novel metastasis-associated lncRNAs and nearby protein coding genes (a) *LOC101448202* (b) *LOC101928307* (c) *LINCR-0002* (d) *LINC00842* (e) *GNAS-AS1*. Analysis of publicly available microarray datasets comparing tumor biopsies of CRC patients without metastasis to liver tumor biopsies of CRC patients with metastasis. (f) *PRKCQ-AS1*. n = 6 per group. *** p <0.001. (g-i) *LINC00635, LINC00842, DLX6-AS1*. n = 9 controls, 6 liver metastases. * p <0.05, ** p <0.01.

Next, the findings in this *in vivo* selection study were compared to data extracted from two prior clinical studies that analyzed differences in lncRNAs between primary colon cancers that do not metastasize compared to CRC-derived liver metastases (7, 8). Of the top 20 altered lncRNAs identified from our study (Figure 1f), 5 had corresponding microarray probes that corresponded to the clinical dataset. Of the 5 lncRNAs with probes in our study, 3 were significantly altered in the clinical metastatic samples (Figure 3f-h): *PRKCQ-AS1* (p<0.001), *LINC00635* (p<0.01) and *LINC00842* (p<0.05). *DLX6-AS1* was altered in a consistent direction to our data though the result was not statistically significant (Figure 3i). These results suggest that the *in vivo*-selected highly metastatic cell line closely modeled clinical observations.

### Down-regulation of lncRNAs identified via *in vivo* selection do not influence *in vitro* metastatic functions

To identify functional assays that could explain the differences in metastatic rates between LS-PAR and LS-3B cells, both cell growth and migration were measured. Over the period of 4 days, cell growth was measured and interestingly the mildly metastatic parental cells (LS-PAR) grew faster than the highly metastatic cells (LS-3B) (Figure 4a). To measure the migratory ability of the cells, an important aspect of metastasis, scratch wound assays were employed (27). Similar to the differences seen *in vivo*, LS-3B cells were able to migrate more efficiently (Figure 4b). Now as a notable metastatic ability, migration can discriminate LS-3B cells compared to LS-PAR cells it was next tested if any lncRNAs identified in the large-scale screen between the highly metastatic cells and their parental counterparts functionally participate in the metastatic ability of migration. Initially these experiments were performed in LS-3B/LS-PAR cells, however these cell lines were unable to be transfected and form monolayers in enough time. Therefore, the highly metastatic SW480 colon adenocarcinoma cells were used to screen lncRNA function. First a subset of the lncRNAs identified in Figure 1d were downregulated with siRNAs, and downregulation was confirmed with RT-PCR (Figure 4c). Next, cells with downregulated lncRNAs were given a scratch wound. The control siRNA treated cells closed 40-70%, depending on the experiment. When compared to the control siRNA treated cells, cells with down-regulated lncRNAs exhibited similar rates of wound closure (Figure 4d). Cells with down-regulated lncRNAs also failed to display differences compared to controls in other functional assays such as invasion and chemotaxis (data not shown). These results suggest that if the lncRNAs identified by *in vivo* metastasis selection do participate in pro-metastatic functions then they must do so through *in vivo* specific mechanisms.

**Figure 4:**
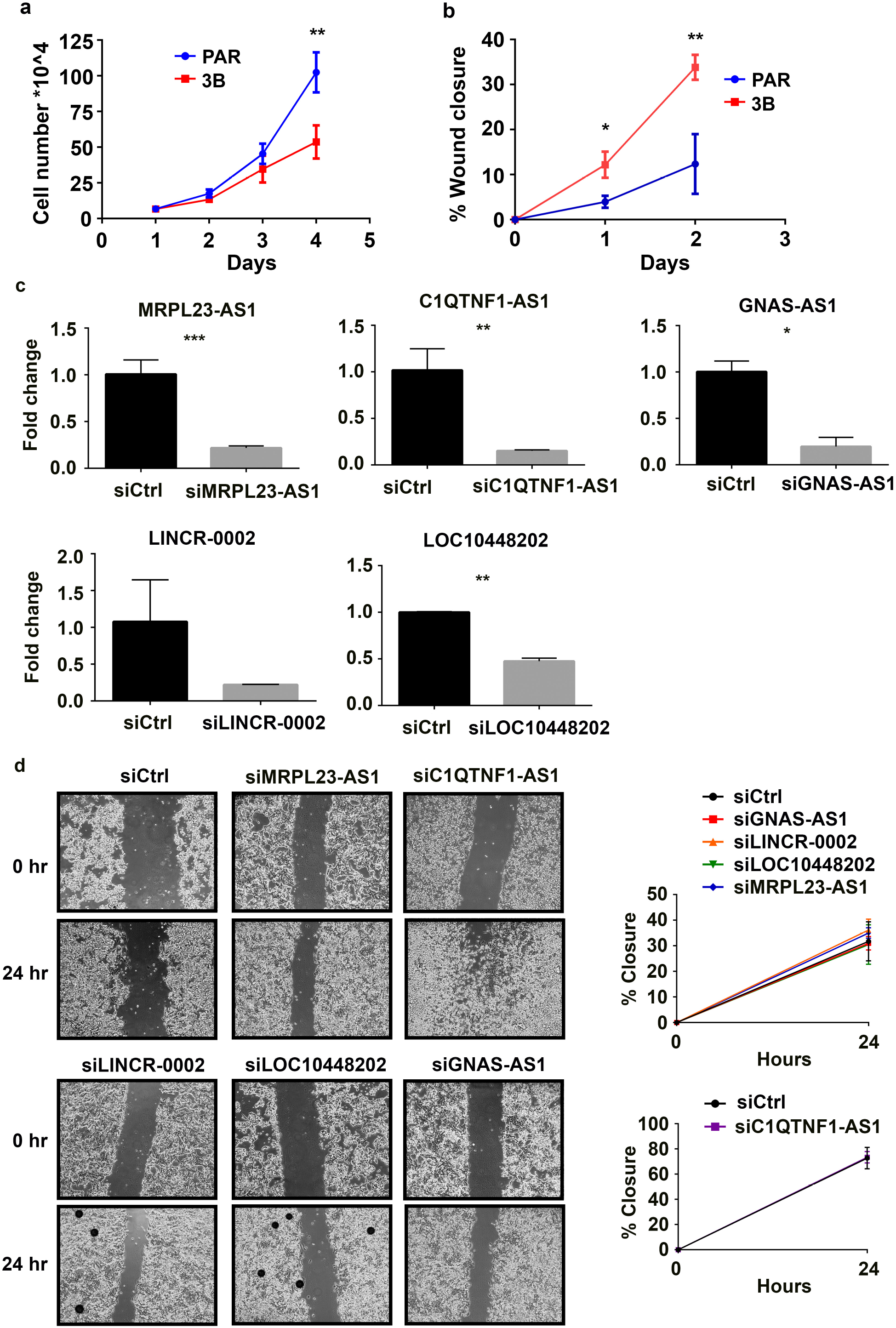
Down-regulation of lncRNAs identified via *in vivo* selection do not influence *in vitro* metastatic functions. (a) Equal numbers of cells were plated and each day after cell numbers were counted. Graph is representative of three separate experiments. (b) Monolayers of cells were wounded and at 24 and 48 hours post-wounding images were taken of the same location and wound closure was measured. Graph is representative of three separate experiments. (c) Real time PCR was used to measure the levels of lncRNAs after 96 hours of siRNA mediated down-regulation in SW480 cells. n= 3 per group. Representative graphs from 3 independent experiments. * p <0.05, ** p <0.01, *** p <0.001. (d) Images were taken immediately after scratch wound and 24 hours later in cells with down-regulated lncRNAs. 3 independent experiments. n= 3 per group.

## Discussion

With the advent of high-throughput analysis of the transcriptome, it has become evident that protein coding genes are not the only players in health and disease. A previous study utilized *in vivo* selection to identify one category of functionally important non-protein coding RNAs, microRNAs, in metastasis (30). The results presented in this manuscript utilize this model to identify and characterize another subset of non-protein coding genes, long non-coding RNAs, in colon to liver metastasis. While lncRNAs have not been profiled in models using *in vivo* selection, functional screens and clinical samples have identified pro-metastatic lncRNAs. The most well-known lncRNA with an association to metastasis that was observed in this study to be altered in *in vivo* selected CRC cells was *H19*, which was 7-fold increased by microarray and 5-fold by RT-PCR in highly metastatic daughter cells. Interestingly, alterations in other well-known metastasis associated lncRNAs, *HOTAIR* and *MALAT1*, were not observed in this model of colon to liver metastasis (43). However, *HOTAIRM1* was observed to be elevated 3.6-fold (FDR < 8.3E-5) in the highly metastatic cells (data not shown) and *HOTAIRM1* has been shown to function similarly to *HOTAIR* in HOX chromatin remodeling (46). The differences in alterations between this study and studies on *MALAT1* and *HOTAIR* could be due to cell type specific differences. Other than *H19*, three other lncRNAs that have been associated with metastasis were identified in this study to be altered in *in vivo* metastasis selected CRC cells: *LINC00842, CASC9, DLX6*-AS1. *LINC00842* has been shown to be TGFB responsive and both *LINC00842* and *CASC9* were previously implicated in tissue invasion (1, 29, 35, 38, 52).

A literature search revealed no studies for several lncRNAs observed to be altered in this model of *in vivo* metastasis selection (*LOC101928201, LOC101448202, PRKCQ-AS1, MRPL23-AS1, LINCR-0002)*. However, SNPs in *LOC101928201* and *LOC101448202* have previously been associated with cancer (53). One clue to the function of these lncRNAs is their genomic surroundings, as lncRNAs have been shown to regulate transcription of nearby genes (36, 37, 45). In figure 3, an analysis of lncRNAs identified to be altered in highly metastatic cells highlighted their proximity to highly pro-metastatic protein coding genes.

In this study, down-regulation of lncRNAs did not affect expression of nearby protein coding genes or result in functional differences measured by *in vitro* metastasis assays. One explanation for this is that pro-metastatic functions may be *in vivo* specific mechanisms or rely on factors that are not adequately modeled in our cell culture system. For example, a caveat to this study was the lack of an immune system in this model of metastasis. Lately this has been overcome by using primary intestinal cell cultures from cancerous mice such as *Apc*^*min*^ mice in *in vivo* metastasis studies instead of human cell lines (12). However, an inherent problem with studying long-non coding RNAs is the lack of conservation in mice.

Another caveat of this study which is the lack of genomic removal to study the function of lncRNAs. This may account for the absence in both changes in expression of neighboring genes and functional differences measured by metastasis assays. Removal of the lncRNA is likely necessary to study cellular as lncRNAs often regulate nearby genes during their transcription (24). The best method to further this study will be using CRISPR to remove the lncRNAs from LS-3B or LS-PAR cells and then perform the in vivo metastasis assay to assess the roles for the lncRNAs identified in colorectal cancer metastasis.

When comparing the lncRNAs identified by *in vivo* selection and clinical samples only 3 associations were found, however the lack of microarray probes for the newly annotated lncRNAs hampered efforts. To better understand if the lncRNAs identified in this study are relevant to disease, future studies using up to date microarrays or a re-analysis of RNA-seq data could be performed. Additionally, this study should be compared to other models of *in vivo* metastasis such as brain or lung to identify lncRNAs common to metastasis.

This study on lncRNAs poses an exciting opportunity for CRC metastasis research as antisense nucleic acid drug inhibitors display much greater specificity compared to current systemic treatments for CRC metastasis (34). New biomarkers and new therapeutic targets can be gleaned from the differentially expressed lncRNAs identified by *in vivo* selection.

---

**Supplemental Table 1.**
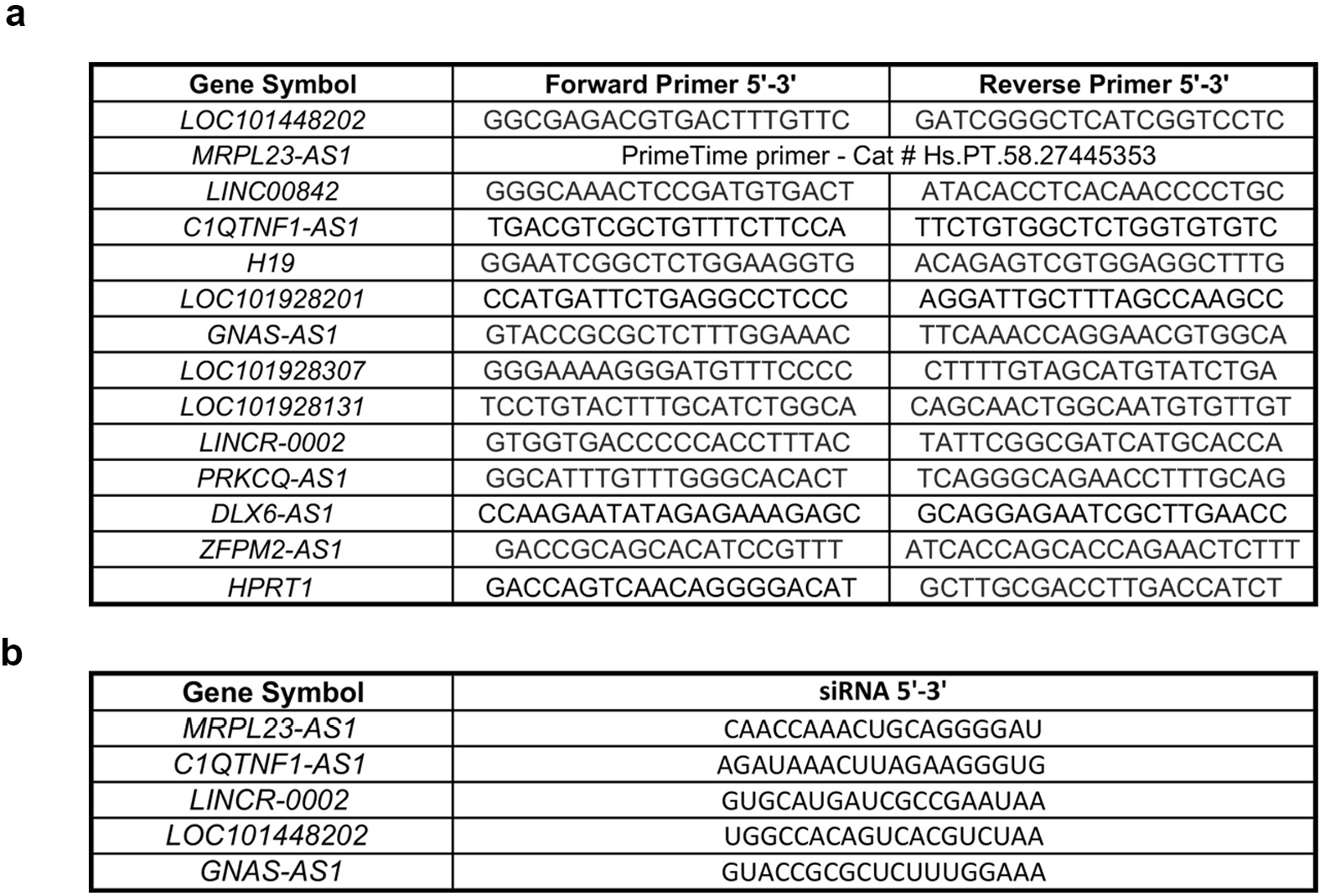

