## Supplemental Table 1 for "*In vivo* selection reveals long non-coding RNAs implicated in colon to liver metastasis"

**a**

| Gene Symbol | Forward Primer 5'-3' | Reverse Primer 5'-3' |
|---|---|---|
| *LOC101448202* | GGCGAGACGTGACTTTGTTC | GATCGGGCTCATCGGTCCTC |
| *MRPL23-AS1* | PrimeTime primer - Cat # Hs.PT.58.27445353 | |
| *LINC00842* | GGGCAAACTCCGATGTGACT | ATACACCTCACAACCCCTGC |
| *C1QTNF1-AS1* | TGACGTCGCTGTTTCTTCCA | TTCTGTGGCTCTGGTGTGTC |
| *H19* | GGAATCGGCTCTGGAAGGTG | ACAGAGTCGTGGAGGCTTTG |
| *LOC101928201* | CCATGATTCTGAGGCCTCCC | AGGATTGCTTTAGCCAAGCC |
| *GNAS-AS1* | GTACCGCGCTCTTTGGAAAC | TTCAAACCAGGAACGTGGCA |
| *LOC101928307* | GGGAAAAGGGATGTTTCCCC | CTTTTGTAGCATGTATCTGA |
| *LOC101928131* | TCCTGTACTTTGCATCTGGCA | CAGCAACTGGCAATGTGTTGT |
| *LINCR-0002* | GTGGTGACCCCCACCTTTAC | TATTCGGCGATCATGCACCA |
| *PRKCQ-AS1* | GGCATTTGTTTGGGCACACT | TCAGGGCAGAACCTTTGCAG |
| *DLX6-AS1* | CCAAGAATATAGAGAAAGAGC | GCAGGAGAATCGCTTGAACC |
| *ZFPM2-AS1* | GACCGCAGCACATCCGTTT | ATCACCAGCACCAGAACTCTTT |
| *HPRT1* | GACCAGTCAACAGGGGACAT | GCTTGCGACCTTGACCATCT |

**b**

| Gene Symbol | siRNA 5'-3' |
|---|---|
| *MRPL23-AS1* | CAACCAAACUGCAGGGGAU |
| *C1QTNF1-AS1* | AGAUAAACUUAGAAGGGUG |
| *LINCR-0002* | GUGCAUGAUCGCCGAAUAA |
| *LOC101448202* | UGGCCACAGUCACGUCUAA |
| *GNAS-AS1* | GUACCGCGCUCUUUGGAAA |
